# Lithium drives coordinated changes in the mouse synaptic phosphoproteome

**DOI:** 10.64898/2026.04.16.718903

**Authors:** Brooke A Prakash, Ishani Shah, Iolanda Vendrell, Roman Fischer, Russell G Foster, Aarti Jagannath, Sridhar R Vasudevan

## Abstract

Lithium is the gold standard mood stabiliser used to treat cycling mania and depression in bipolar disorder. Despite seven decades of clinical use, the mechanisms of its mood stabilisation are incompletely understood, fundamentally limiting development of improved alternatives. Two established lithium targets, glycogen synthase kinase 3β (GSK3β) and inositol monophosphatase, both modulate phosphorylation, suggesting lithium may exert broad effects on neuronal phosphorylation networks. We performed a discovery-phase *in vitro* screen of 140 kinases at 10mM LiCl and demonstrated that lithium inhibits 17 kinases beyond GSK3β. We therefore used untargeted quantitative phosphoproteomics to create a comprehensive map of lithium’s neural phosphorylation signature in lithium-treated mouse synaptoneurosomes. Samples were collected at dawn and dusk to match the peaks in phosphorylation that are induced by the sleep/wake cycle. Pathway analysis revealed convergence on synaptic plasticity, neurotransmitter release, and chemical transmission. Critically, lithium-sensitive phosphoproteins are significantly enriched in bipolar disorder genome-wide association study (GWAS) loci, providing independent genomic evidence that the phosphorylation networks we identified are relevant to bipolar pathophysiology. We identified novel kinase targets and phosphorylation sites not previously associated with lithium’s mechanism of action and tied them to bipolar pathology. We further refined existing models of lithium’s action by showing that GSK3β inhibition is temporally restricted to dawn, indicating cross talk with sleep/wake cycles of phosphorylation. Overall, our data demonstrate that lithium’s pleiotropic effects result from coordinated multi-kinase network reorganisation rather than single-target inhibition — a principle with direct implications for rational polypharmacology in mood stabiliser development.

## Introduction

Bipolar disorder (BD) is a manic-depressive mood disorder that affects 1%-5% of the world population^1^. There are very few medications that can treat both the mania and depression of BD; lithium is the notable exception. Nevertheless, despite seven decades of clinical use, the molecular mechanisms underlying lithium’s mood-stabilising effects remain poorly defined. This fundamental knowledge gap has impeded the development of alternative treatments that could retain lithium’s therapeutic efficacy whilst avoiding its narrow therapeutic window and dose-limiting side effects that contribute to patient noncompliance and discontinuation of treatment^2^.

Two molecular targets of lithium have historically dominated research into its therapeutic efficacy – the kinase glycogen synthase kinase 3 β (GSK3β) and the phosphatase inositol monophosphatase (IMPase). Lithium inhibits GSK3β directly, through competition with magnesium during phosphate transfer, and indirectly through activation of upstream pathways^3–5^. GSK3β inhibition has neuroprotective and mood-altering properties and has therefore been proposed to underlie lithium’s therapeutic effects^6, 7^. Lithium also uncompetitively inhibits IMPase, reducing inositol supply and neuronal signalling^8, 9^. This observation led to the inositol depletion hypothesis, which suggests that lithium reduces aberrant signalling during mania through reducing neuronal inositol supplies^10, 11^. GSK3β inhibition and inositol depletion do not phenocopy all of lithium’s cellular effects, however, pointing to a broader point of regulation by lithium.

Given that lithium has been shown to inhibit GSK3β via magnesium competition, it is plausible that lithium could broadly inhibit other kinases as well. Previous studies have documented lithium’s effects on additional kinases in isolated experimental contexts, revealing substrate-specific alterations. Activated c-Jun NH2-terminal kinases were detected in the frontal cortex and hippocampus of lithium-treated male rats while lithium’s alterations of protein kinase (PK) C and A activity were shown to be substrate-specific in the hippocampus^12, 13^. Primary cultures of rat cerebellar granule cells have further demonstrated a lithium-driven increase in Akt activity^3^. Work on male bipolar patient-derived iPCSs revealed alterations in MAPK, PKC, GSK3, PKA, AMPK and Akt-driven phosphorylation in lithium-responsive cells. Further alterations occurred upon lithium treatment, but it was also noted that the directionality of these changes is substrate-specific^14^.

We hypothesised that lithium induces widespread reorganisation of synaptic phosphorylation networks beyond its canonical targets, engaging multiple kinase families to produce its pleiotropic effects. Given that protein phosphorylation enables rapid, reversible modulation of synaptic function and that synaptic dysfunction is implicated in bipolar disorder pathophysiology, comprehensive characterisation of lithium-induced phosphorylation dynamics could explain how this single drug simultaneously modulates the diverse neuronal processes underlying mood stabilisation.

Emerging evidence from complex therapeutic contexts suggests that efficacy in robust biological systems frequently emerges from coordinated multi-target modulation rather than selective inhibition of individual components. In oncology, phosphoproteomic analyses of kinase inhibitor resistance have demonstrated that single-target agents fail through network rewiring, with alternative kinases compensating for inhibited targets – a finding that has redirected therapeutic strategies towards approaches targeting multiple network nodes simultaneously^15^. Similarly, physiological state transitions including sleep-wake cycles involve coordinated reorganisation of synaptic phosphorylation networks that cannot be recapitulated by modulating individual kinases^16^. These precedents raise the possibility that lithium’s distinctive therapeutic profile, which has resisted explanation through single-target models despite decades of investigation, reflects an analogous network-level pharmacology.

Here, we performed an *in vitro* assay of kinase activity during 10mM LiCl administration to confirm the existence of lithium-sensitive kinases beyond GSK3β. We further performed quantitative phosphoproteomics of synaptoneurosomes isolated from mouse forebrain following chronic administration of lithium at therapeutically relevant serum concentrations. We analysed male and female mice separately at dawn and dusk timepoints to capture temporal and sex-specific dynamics of synaptic phosphorylation. Our findings provide a comprehensive map of lithium’s synaptic phosphorylation network and identify previously unrecognised molecular targets for mood stabiliser development.

## Methods

### *In vitro* Screening of Kinase Activity

The kinase activity screen was performed as previously described at the MRC Protein Phosphorylation and Ubiquitylation Unit International Centre for Kinase Profiling^17^. Briefly, each kinase was diluted so that its activity was linear with respect to time and enzyme concentration. Protein substrate and diluted kinase were incubated in concentrated assay buffer (50mM Tris-HCl, 0.1mM EGTA and 10mM magnesium acetate with additional cofactors as needed for each kinase) for 5 minutes. The protein kinase reaction was initiated at room temperature with the addition of [γ-33P]ATP at or below the Km of ATP for each kinase. The reaction was stopped with 0.5 M orthophosphoric acid and each reaction mixture was spotted onto a P81 filter plate (Whatmann) using a unifilter harvester (PerkinElmer). Scintillation fluid was added and radioactivity of the protein substrates, blank filter plates and ATP alone was measured with a TopCount NXT scintillation counter (PerkinElmer). Each plate was compared to quality controls assessing free ATP contamination and autophosphorylation of each kinase. Each reaction was carried out in duplicate.

### *in vivo* Drug administration

8-week-old wild-type C57BL/6J (Charles River Laboratories) male and female mice were individually housed on running wheels under 12:12 LD cycles for one week. 7mM LiCl (Sigma-Aldrich) was dissolved in drinking water and administered *ad libitum* for one week to gradually build up the dose, reducing adverse effects^18^. A full dose of 14mM LiCl was then administered *ad libitum* for two weeks before sample collection^19^. A second bottle of 0.9% saline (NaCl - Sigma-Aldrich) was provided throughout lithium treatment to mitigate ion imbalances and toxic side effects^20, 21^. Vehicle mice received plain drinking water.

### Synaptoneurosome Isolation

Dusk samples were collected at ZT11 and dawn samples were collected at ZT21 to match the peaks of cycling phosphorylation in the synaptoneurosomes of male mice *in vivo*^16^. Synaptoneurosomes were isolated using Percoll gradients as previously described^22^. Briefly, animals were sacrificed with cervical dislocation and the whole brain was extracted. The brain was then homogenised in a Dounce homogeniser in ice cold isotonic sucrose buffer (0.32M Sucrose, 1mM EDTA, 5mM Tris, pH 7.4) before being centrifuged at 1000g for 10 minutes. The resulting supernatant was loaded over layered Percoll (Cytiva) gradients (3%, 10%, 15% and 23%) and spin at 31,000g for 5 minutes. All fractions between 10% and 23% were collected, diluted to ~30mLs in sucrose buffer and spun at 20,000g for 30 minutes. The resulting pellet containing synaptoneurosomes was resuspended in 5mM Tris-HCl, pH 7.4 (Sigma-Aldrich) and flash frozen for phosphopeptide enrichment. Throughout synaptosome isolation, all centrifugation steps were performed at 4ºC and all solutions were kept ice cold and were supplemented with 50uM DTT (Sigma-Aldrich), 0.1mM PMSF (Sigma-Aldrich), 1X cOmplete EDTA-free Protease Inhibitor cocktail (Roche), and 1X Halt^™^ Phosphatase Inhibitor cocktail (Thermo Scientific).

### Protein digest and Phosphopeptide Enrichment

Synaptosome extracts were lysed in RIPA buffer (50mm Tris-HCl, 150mM NaCl, 0.1 SDS, 0.5% sodium deoxycholate, 1% NP40) supplemented with protease and phosphatase inhibitors. Samples were digested and phosphoenriched as previously described^23^. Briefly, proteins were precipitated using methanol:chloroform to remove the SDS, solubilised in 8M urea in HEPES at pH8 and digested with trypsin (1:50 trypsin:protein ratio) overnight at 37ºC. Desalted peptides were subjected to a phosphopeptide enrichment. This was performed using the default Agilent Bravo Assaymap liquid handler workflow for titanium dioxide (TiO2) cartridges. Enriched phosphopeptides samples were dried down and resuspended in 2% acetonitrile, 0.1% TFA.

### LC-MS/MS and Data Processing

Phosphopeptide enriched samples were analysed by LC-MS/MS using the U3000 nUHPLC connected to the Orbitrap Fusion Lumos (male set) or the Vanquish Neo UHPLC connected to an Orbitrap Ascend (female set).

For the male samples, peptides were trap onto a PepMapC18 trap column (300µm x 5mm, 5µm particle size, Thermo Fisher) and separated on a 50cm EasySpray column (ES803, Thermo Fisher) using a 60-minute linear gradient from 2 % to 35% buffer B (A: 5% DMSO, 0.1% formic acid; B: 5% DMSO, 0.1% formic acid in acetonitrile) at 250 nl/min flow rate. Data were acquired in the Orbitrap Fusion Lumos in data dependent mode (DDA) with the advance peak detection (APD) switched on (as described https://doi.org/10.1016/j.celrep.2024.114152). Full scans were acquired in the Orbitrap at 120 k resolution over a m/z range 400-1500, AGC target of 4e5 and S-lens RF of 30. MS2 scans were obtained in the Ion trap (rapid scan mode) with a Quad isolation window of 1.6, 40% AGC target and a maximum injection time of 35 ms, with HCD activation and 28% collision energy.

For the female samples, peptides were analysed using the Vanquish Neo operated in “Trap and Elute” mode using a PepMap Neo trap (5 μm, 300 μm x 5 mm; Thermo Fisher) with backflash and EASY-SPRAY PepMapNeo column (50 cm x 75 um, 1500 bar; Thermo Fisher). Tryptic peptides were trapped and separated over a 75 min gradient, going from 3 to 20% B (0.1% FA ACN) in 45 min, to 35% in 15 min, up to 99% B in 1 min and then staying at 99% for 14 min. The flow rate was maintained at 300 nL/min throughout the gradient. MS data were acquired in data dependent mode (DDA). MS1 scans were collected in the Orbitrap at a 120K at m/z 200 over m/z range of m/z 380-1500. The MS1 normalised AGC was set at 100% with a maximum injection time set to auto and a RF lens to 30%. The top 20 most intense ions were then acquired in the Orbitrap at 15K resolution, with a normalised AGC target of 100%, maximum injection of 27 ms and a HDC collision energy of 30%.

MS raw files were analysed using MaxQuant (v1.6.14 for the male set and v2.3.0.0 for the female set). Files were searched against the UniProt-Swissprot mouse database (Feb2021 for the male set and Oct 2023 for the female) using the in-build Andromeda data-search engine. Trypsin was selected as enzyme (up to 2 missed cleavages), carboamidomethylation (C) as fixed modification and Deamidated (NQ), Oxidation (M) and phosphorylation (STY) as variable modifications. Protein false discovery rate was set up at 1%. Data were quantified using the label free quantitation (LFQ) with match between runs (MBR) enabled.

### Data Analysis and Visualisation

Phosphoproteomic data was analysed with Phosphomatics^24^. Phosphosites detected in fewer than 2 samples in any treatment group were excluded, then missing data was imputed from the treatment group minimum. The resulting phosphorylation data was log2 transformed and median normalised. Principal component analysis, ANOVA, and Kinase-Substrate Enrichment Analysis (KSEA) were all performed with Phosphomatics’ inbuilt functions. The transformed and normalised data was analysed with ANOVA followed by Benjamini-Hochberg corrections to select significant phosphopeptides (PAdj≤0.2). KSEA was performed to identify kinases that are associated with the phosphorylation sites that differ in abundance between treatment groups. Kinases with at least two identified substrates with a NetworKIN score of 4, P≤ 0.1, and predicted fold change in activity ≥15% were considered significant going forward.

Fisher’s exact test was performed to assess whether the genes of lithium-sensitive phosphoproteins were significantly enriched among BD GWAS-implicated genes (using Open Targets v25.12 GWAS credible set threshold > 0.1, P ≤ 0.05)^25^. BD GWAS-implicated genes were compared with the Kinase Library 2024 in Enrichr using a Fisher’s exact test to predict which kinases have substrates in BD-related genes (PAdj≤0.05)^26^.

Volcano, bar, lollipop and PCA plots were produced using GraphPad Prism (10.4.2) or R Studio (2024.04.2+764). Functional enrichment analysis for Gene Ontology Biological Processes (GO:BP) and Cellular Components (GO:CC) was performed using the STRING database (version 12) with FDR≤ 0.05^27^. Cytoscape (3.10.4) was used to visualise biomolecular interactions with a confidence score ≥ 0.4 between hit phosphopeptides^28^. Visualisation of the kinome tree was performed with KinMap^29^ and pathway/neuron visuals were created in BioRender. Fig. 2A created in BioRender. Prakash, B. (2026) https://BioRender.com/k8iihia. Fig. 5C created in BioRender. Prakash, B. (2026) https://BioRender.com/5l7tq3l.

## Results

To examine the extent of lithium’s kinase inhibition *in vitro*, we screened the activity of 140 kinases in the presence of 10mM LiCl, a concentration consistent with established kinase profiling protocols and previously used to characterise lithium’s kinase selectivity profile. Lithium inhibited the activity of 18 kinases including GSK3β greater than 20% (Fig. 1A). These kinases fell across the kinase phylogenetic tree, suggesting that inhibition of activity by lithium is not specific to a particular kinase structure (Fig. 1B).

**Figure 1:**
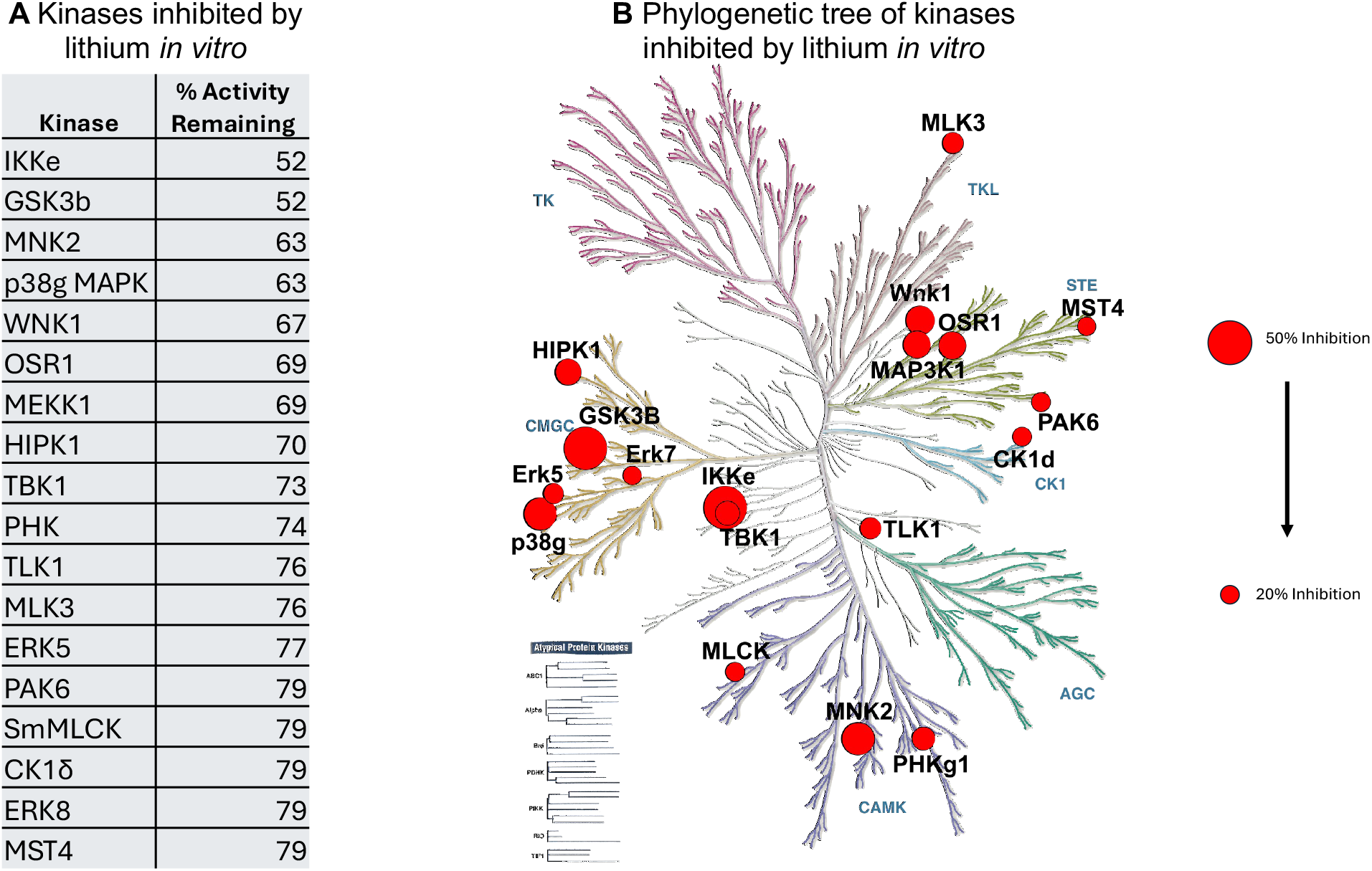
Lithium inhibits 18 kinases *in vitro*. **A** *in vitro* assay of kinase activity during 10mM LiCl treatment. **B** Affected kinases are plotted on the kinase phylogenetic tree.

Having demonstrated *in vitro* that lithium suppresses a network of 18 kinases, we next sought to determine if this multi-kinase modulation translates *in vivo*. To test this, we performed quantitative phosphoproteomics on synaptoneurosomes isolated from lithium-treated mice. We administered 14mM LiCl in the drinking water of male mice for 2 weeks following a paradigm known to produce therapeutic serum lithium levels for bipolar disorder^19^. Synaptoneurosomes were isolated from brain homogenate collected from these lithium and vehicle-treated mice at dawn (ZT21) and dusk (ZT11) (Fig. 2A). These timepoints were chosen based on previous work demonstrating that major phosphorylation events occur in the brain twice daily at dawn and dusk^16^.

**Figure 2:**
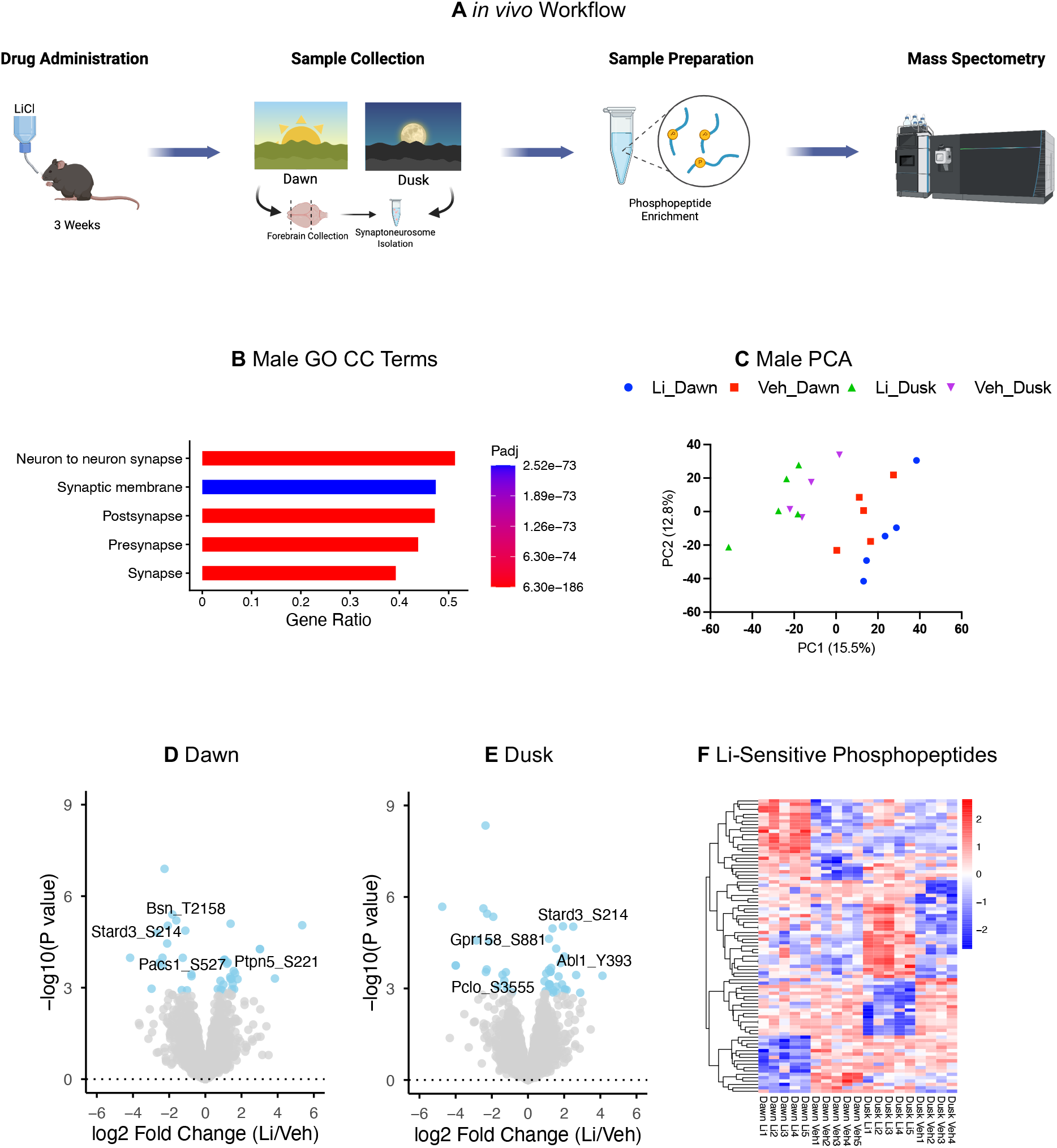
Lithium treatment induces time-of-day specific reorganisation of the synaptic phosphoproteome. **A** *in vivo* experimental workflow. **B** Quality control validation showing Uniprot keyword enrichment for synapse-related words among phosphopeptides detected in male mice. **C** PCA analysis of phosphopeptides detected in male mice (N=4-5 per timepoint). Volcano plots show lithium-sensitive phosphosites at **D** dawn and **E** dusk in male mice (N=4-5 per timepoint). **F** Heatmap showing distinct temporal organisation of lithium-responsive phosphosites in males.

Quantitative mass spectrometry identified 6657 phosphosites in 2018 proteins in the synaptoneurosomes. Gene Ontology (GO) enrichment analysis of cellular components (CC) confirmed the integrity of synaptoneurosome preparation with cellular components such as “synapse” and “synaptic membrane” significantly enriched in the phosphopeptides detected across all samples (Fig. 2B). As expected, principal component analysis (PCA) showed a strong time-dependent separation of samples (Fig. 2C). 87 of the detected phosphopeptides were specifically lithium-sensitive (Fig. 2D-F). Each of these phosphopeoptides were time-of-day specific except for Stard3 S214, which was decreased by lithium at dawn and increased by lithium at dusk (Fig. 2D,E). The limited overlap between dawn and dusk timepoints indicate that lithium’s synaptic effects are not constitutive, but rather gated by time-of-day-dependent molecular states.

Analysis of the proteins with lithium-sensitive phosphorylation sites revealed an interconnected network of proteins involved in synaptic organisation and chemical transmission (Fig. 3A,B). This network was less extensive at dawn, as only 6 biological processes were significantly altered (Fig. 3C). Nevertheless, lithium did notably increase inhibitory phosphosite S221on phosphatase PTPN5/STEP61 – a site whose inhibition has been associated with synaptic strengthening in prior studies^30^. At dusk, lithium altered phosphorylation in wide-ranging processes involved in synaptic growth, signalling and organisation (Fig. 3D). This included activation of Abelson non-receptor tyrosine kinase 1 (Abl1) at Y393, a site previously linked to reduced spontaneous postsynaptic currents^31, 32^. Lithium also increased Gpr158 S881, a site which is increased by treatment with D1R agonist SKF81297^33^.

**Figure 3:**
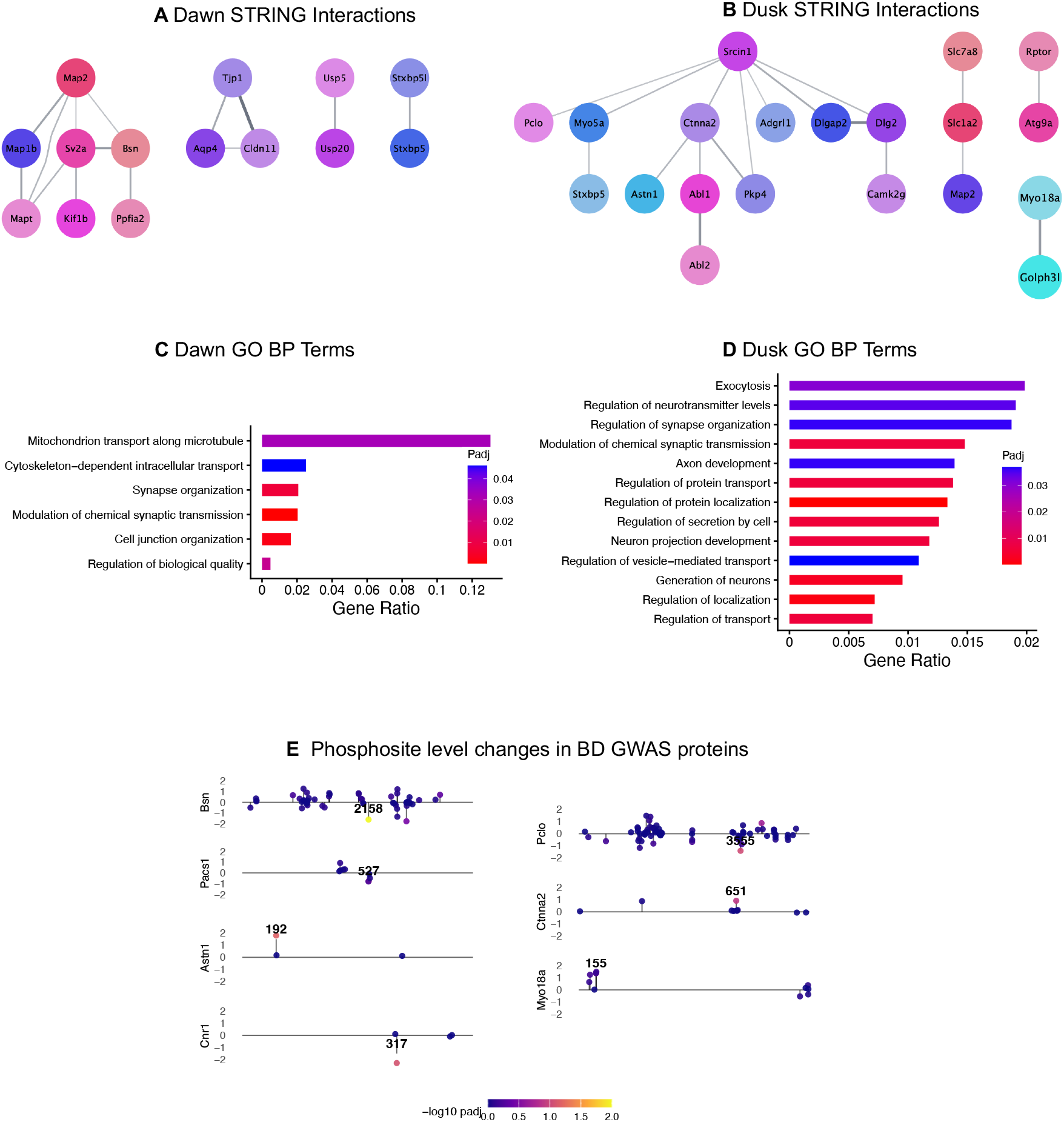
Lithium alters the phosphorylation of proteins involved in synaptic organisation and transmission. STRING analysis was used to visualise protein-protein interaction networks among the lithium-sensitive peptides on hits from **A** dawn **B** dusk. **C** All GO BP enriched terms from dawn lithium-sensitive phosphosites. **D** Representative GO BP terms enriched at dusk. **E** Lollipop plots of phosphosite level changes on the backbone (X-axis) of proteins with a BD GWAS score >0.1, colours indicative of adj p-value (scale below) and direction and length of stick indicative of fold change.

To assess the therapeutic relevance of the lithium-sensitive phosphoproteins, we compared them with genes that have been linked to BD in GWAS on the Open Targets platform^25^. Fisher’s exact test demonstrated that 7 genes with lithium-sensitive phosphosites (PAdj ≤ 0.2) were significantly enriched among BD GWAS-implicated genes (OR = 2.27, p = 0.05) (Fig. 3E). These proteins play a role in regulating presynaptic vesicle organisation, endocannabinoid signalling, neurotransmitter release, and cell adhesion.

We further performed an independent replication in female mice to see if there was pathway-level convergence in lithium’s alteration of the synaptic phosphoproteome between sexes. Quantitative mass spectrometry identified 2816 phosphosites in 1211 proteins in the female synaptoneurosomes. GO CC enrichment analysis of these proteins once again indicated the integrity of synaptoneurosome preparation (Fig. 4A). 83.98% of phosphopeptides detected in female synaptoneurosomes were also detected in male synaptoneurosomes showing robust overlap between the two datasets.

**Figure 4:**
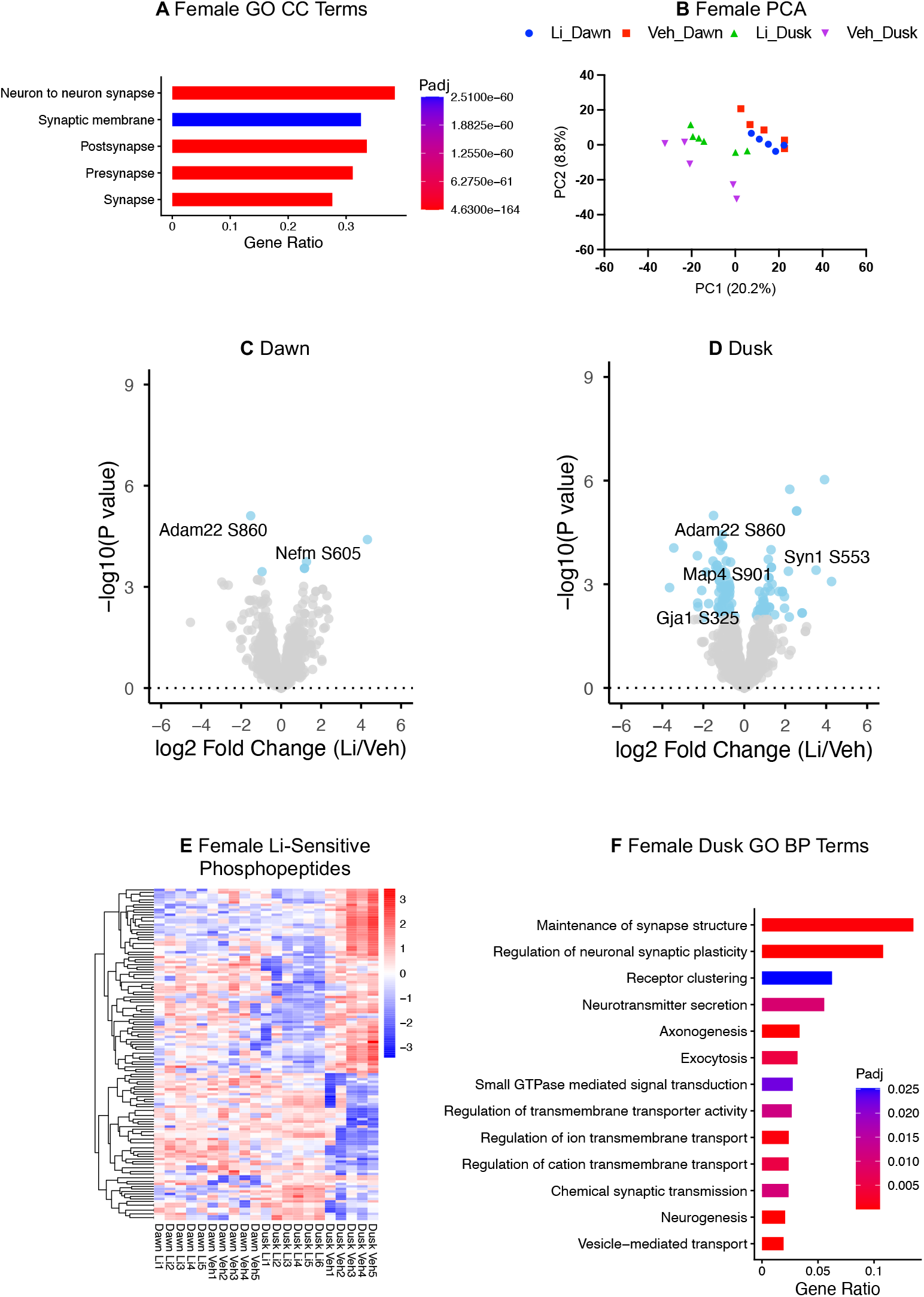
Lithium treatment also induces time-of-day specific reorganisation of the female synaptic phosphoproteome. **A** Quality control validation showing Uniprot keyword enrichment for synapse-related words among phosphopeptides detected in female mice. **B** PCA analysis of phosphopeptides detected in female mice (N=5-6 per timepoint). Volcano plots showing lithium-sensitive phosphosites at **C** dawn and **D** dusk in female mice (N=5-6 per timepoint). **E** Heatmap showing distinct temporal organisation of lithium-responsive phosphosites in females. **F** STRING analysis was used to visualise protein-protein interaction networks among the lithium-sensitive peptides on hits from dusk in female synaptoneurosomes.

As with the males, there was a strong time-of-day effect on synaptic phosphorylation (Fig. 4B) and each of the 133 lithium-sensitive phosphosites were time-of-day specific, except Adam22 S860 which was reduced at dawn and dusk (Fig. 4C-E). At dawn, lithium only altered 6 phosphosites, mirroring the lack of a network-coordinated lithium response in dawn in males (Fig. 3C). At dusk, lithium-sensitive phosphopeptides in females also formed a highly interconnected network with phosphosites changing in proteins that regulate synaptic structure, receptor localisation and signalling, closely mirroring the network changes in males (Fig. 4F).

As lithium altered the phosphoproteome in male and female mice, we performed kinase-substrate enrichment analysis (KSEA) to ascertain which kinases are lithium-sensitive. Across timepoints, lithium significantly altered the substrates of 29 kinases (Fig. 5A,B).

**Figure 5:**
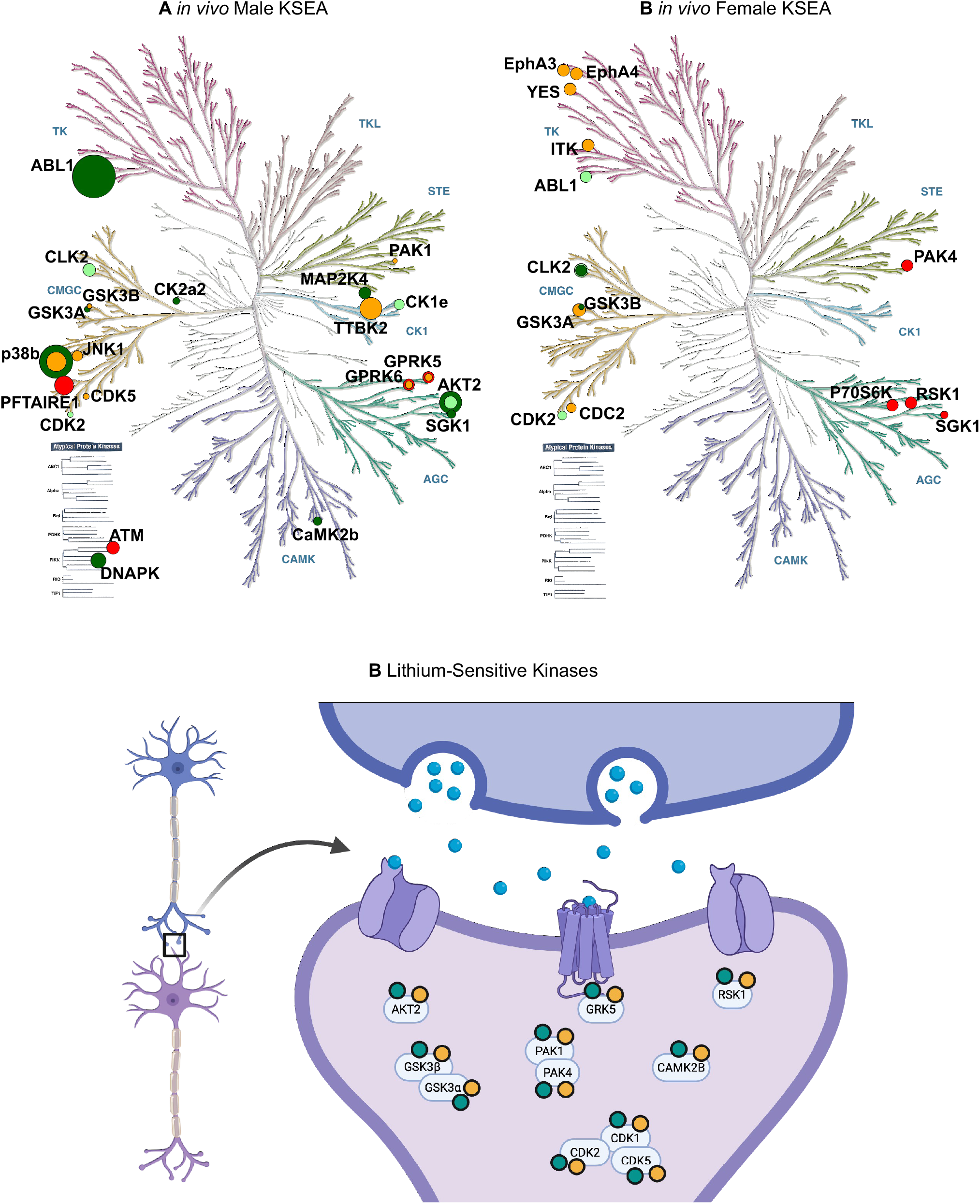
Lithium-Sensitive Kinases are Implicated in the Pathophysiology of Bipolar Disorder. The phylogenetic relatedness of lithium-sensitive kinases in **A** male synaptoneurosomes and **B** female synaptoneurosomes. Yellow denotes increased activity at dawn, orange denotes reduced activity at dawn, green denotes increased activity at dusk, and red denotes reduced activity at dusk. Dot size correlated to fold change of activity. **C** Schematic representation of lithium-sensitive kinases with multiple lines of evidence. Green denotes *in vivo* evidence from our phosphoproteomics, yellow denotes evidence from bipolar GWAS.

The kinases consistently predicted to drive lithium’s phosphorylation signature are involved in Wnt, G protein-coupled receptor (GPCR), and stress/inflammatory signalling, suggesting an overarching role for lithium as a synaptic signalling modulator. Supporting previous literature, the lithium-sensitive phosphoproteome indicated an increase in Wnt signalling via activation of AKT2 and inhibition of GSK3β (males) and GSK3α (females) at dawn. This GSK3β inhibition was temporally specific as its substrates were significantly increased in males and females at dusk, a nuance invisible to *in vitro* assays.

Lithium treatment also induced phosphoproteomic changes in effectors of GPCR signalling. Substrates of GPCR kinase (GRK) 5 and 6 were significantly reduced in males by 36.42% at dusk and 23.63% at dawn. Lithium-treated females mirrored this trend as GRK5 and 6 substrates were also trending towards a 23.54% reduction at dusk (p=0.16). GRK5 and 6 promote GPCR desensitisation, thus their inhibition would prolong GPCR signalling at the synapse^34^. Strikingly, GRK5 and GRK6 emerged as the most consistently inhibited kinases across both sexes and timepoints in vivo.

Reduction of activity in JNK1, RSK1 and PAK1/4 pointed to reduced stress and inflammatory signalling at dawn and dusk, except for p38β (MAPK11), whose substrates were increased by lithium at dusk and reduced at dawn in males.

Finally, we explored whether there was a link between lithium-sensitive kinases and BD pathophysiology. To estimate kinases that are dysregulated in BD, we input BD GWAS-implicated genes (threshold > 0.1) into the 2024 Kinase Library to see which kinases had substrate proteins significantly enriched in the BD GWAS gene set (PAdj≤ 0.05)^35, 36^. 11 of these BD-related kinases were also altered by lithium *in vivo* (Table 1), suggesting broad overlap between lithium-sensitive kinases and BD pathophysiology, including GRK5 (Fig. 5B).

**Table 1:** Lithium-sensitive Kinases with substrates significantly enriched in BD GWAS.

| Kinase | Substrate Overlap | Padj | Odds Ratio | Synaptic BD GWAS Overlap | Synaptic Hits/ Total Hits | Synaptic BD GWAS Genes |
| --- | --- | --- | --- | --- | --- | --- |
| PAK1 | 34/795 | 0.009 | 1.80 | 11 | 0.32 | HTT; CACNA1C; SYNE1; PCLO; DRD2; PAK5; PLCL1; ATP2B2; ANK3; DLG5; DOCK2 |
| CAMK2B | 29/742 | 0.034 | 1.63 | 9 | 0.31 | CTNND1; HTT; ELFN1; RIMS1; GRIN2A; BRAF; BLTP1; CACNB2; DLG5 |
| CDK5 | 30/718 | 0.017 | 1.75 | 9 | 0.30 | HTT; ADD3; PCLO; KCNB1; BRAF; BLTP1; DLG5; SHANK2; SNTB1 |
| P90RSK | 36/787 | 0.003 | 1.94 | 10 | 0.28 | CTNND1; CACNA1C; ELFN1; GRM5; PCLO; BSN; PLCL1; BRAF; BLTP1; DLG5 |
| CDK1 | 29/713 | 0.024 | 1.70 | 8 | 0.28 | HTT; ADD3; PCLO; DRD2; KCNB1; BRAF; BLTP1; DLG5 |
| CDK2 | 34/692 | 0.002 | 2.09 | 9 | 0.26 | ZMYND8; HTT; ADD3; PCLO; DRD2; KCNB1; BRAF; BLTP1; KCNQ5 |
| AKT2 | 35/866 | 0.016 | 1.69 | 7 | 0.20 | HTT; CACNA1C; PCLO; BSN; PLCL1; BRAF; ANK3 |
| GSK3B | 30/740 | 0.023 | 1.69 | 6 | 0.20 | TSHZ3; CACNA1C; SYNE1; PCLO; SLC12A5; ANK3 |
| PAK4 | 42/786 | 0.001 | 2.32 | 8 | 0.19 | ADD3; ELFN1; DRD2; PAK5; SLC12A5; BRAF; ATP2B2; DLG5 |
| GSK3A | 32/712 | 0.007 | 1.90 | 6 | 0.19 | PCLO; CTNNA2; BSN; SLC12A5; BLTP1; DLG5 |
| GRK5 | 27/708 | 0.049 | 1.58 | 5 | 0.19 | CTNND1; PCLO; ANK3; CNKSR2; HNRNPD |

## Discussion

Our comprehensive phosphoproteomic analysis demonstrates that lithium reorganises multiple interconnected kinase networks rather than acting through sustained inhibition of individual targets — a finding consistent with the emerging principle, established in oncology and other complex therapeutic contexts, that robust biological efficacy frequently requires coordinated multi-target modulation rather than selective single-kinase inhibition. Whilst GSK3β and IMPase have been well-established as lithium targets, we identified phosphorylation changes spanning the kinase phylogenetic tree, with affected proteins converging on synaptic organisation, neurotransmitter release, and chemical transmission. Critically, the independent convergence of lithium-sensitive phosphoproteins with BD GWAS loci provides orthogonal genomic evidence that these network-level changes are relevant to bipolar disorder pathophysiology. This coordinated network reorganisation provides a molecular framework for understanding lithium’s pleiotropic effects that fundamentally extends beyond canonical single-target models.

Our phosphoproteomic analysis revealed striking temporal dynamics in lithium’s effects on kinase networks, with the majority of phosphorylation changes showing time-of-day specificity. This temporal organisation challenges simplified models of constitutive target inhibition and suggests that lithium’s effects are integrated within the brain’s endogenous sleep-wake cycles. The temporal patterning of lithium-responsive phosphosites indicates that target engagement is not static but dynamically regulated across the sleep-wake transition. Indeed, the temporal restriction of GSK3β inhibition— occurring at dawn but absent at dusk—suggests that single-target drug design strategies may not fully recapitulate lithium’s coordinated network effects and chronopharmacology. Though contrary to established rhetoric on lithium’s mechanism of action, this finding is not unprecedented as studies of male lithium-responsive iPSCs found that three of the five GSK3β substrates detected were increased after lithium treatment^14^. Rather than canonical targets alone explaining lithium’s cellular actions, our data suggest these targets initiate cascading effects through multiple kinase pathways that collectively modulate synaptic function.

One consistent effect of lithium across timepoints was the inhibition of GRK5 and 6. GRKs drive the desensitization of GPCRs by phosphorylating them to promote β-arrestin recruitment and receptor internalisation. Given that multiple neurotransmitter systems implicated in mood regulation—including dopaminergic, serotonergic, and adrenergic pathways—signal through GPCRs, lithium-mediated modulation of GRK5 and 6 could influence receptor sensitivity and downstream signalling dynamics across these systems. PIP_2_ interacts with GRK4, 5 and 6, specifically, helping them to associate with their target GPCRs^37^. GRK6 in particular has been linked to the desensitisation of dopamine receptor D_2_ and GRK6 knockout mice are hypersensitive to dopamine agonists^38^. The role of dopamine in BD is complicated with hyperdopaminergia hypothesised to underlie mania and dopamine deficiency to underlie depression^39^. It’s plausible that a consequence of PIP_2_ depletion is sensitisation to dopamine which could underlie some of lithium’s antidepressant effects. This represents the first identification of GRK5 and 6 as lithium targets and warrants functional validation in future studies.

This study provides the first comprehensive phosphoproteomic characterisation of lithium’s effects on synaptic signalling networks. Several considerations inform interpretation of these findings. First, our analysis was conducted in mouse synaptoneurosomes rather than human tissue, though the core kinase pathways and synaptic proteins affected by lithium are highly conserved across mammalian species. Second, kinase predictions from KSEA rely on computational analysis of phosphorylation motifs rather than direct kinase activity measurements; however, the concordance between KSEA predictions and the enrichment of KSEA-predicted substrates in BD GWAS loci supports the biological relevance of the computational predictions.

While our findings are anchored in bipolar disorder genetics, the broad synaptic reorganisation we observe has significant implications for neurodegenerative conditions. Emerging evidence highlights lithium’s potential efficacy in Alzheimer’s disease and tauopathies, often attributed to the canonical GSK3β-Tau axis. Our temporal map refines this by demonstrating that GSK3β inhibition is dawn-restricted, suggesting that chronotherapy may be relevant for optimising lithium’s neuroprotective dosing. Furthermore, our dataset reveals lithium-induced modulation of targets implicated in neurodegenerative synaptic depression, notably the inhibitory phosphorylation of STEP61 (PTPN5) at S221. Consequently, this multi-kinase framework may provide a mechanistic foundation for understanding lithium’s therapeutic potential beyond mood stabilisation.

Our findings establish a foundation for future mechanistic studies. Critical questions include identifying whether these phosphorylation events are functionally necessary for lithium’s cellular effects, whether targeting key network nodes could recapitulate lithium’s effects through rational polypharmacology, whether lithium-responsive phosphorylation networks differ between treatment responders and non-responders, and establishing direct links between the molecular signatures we identified and behavioural outcomes in animal models. Investigation of these phosphorylation networks in patient-derived neurones and clinical samples could further inform translation of our molecular findings into improved therapeutic strategies for bipolar disorder.

## Acknowledgements

The authors thank the University of Oxford Biomedical Services for animal husbandry support. This work was supported by the BBSRC (grant BB/N001664/1) and a BMS-Celgene fellowship (2022–2025). The authors declare no conflicts of interest.

## References

1. Akiskal HS, Bourgeois ML, Angst J, Post R, Moller H, Hirschfeld R. Re-evaluating the prevalence of and diagnostic composition within the broad clinical spectrum of bipolar disorders. J Affect Disord 2000; 59 Suppl 1: S5–S30.

2. McKnight RF, Adida M, Budge K, Stockton S, Goodwin GM, Geddes JR. Lithium toxicity profile: a systematic review and meta-analysis. Lancet 2012; 379(9817): 721–728.

3. Chalecka-Franaszek E, Chuang DM. Lithium activates the serine/threonine kinase Akt-1 and suppresses glutamate-induced inhibition of Akt-1 activity in neurons. Proc Natl Acad Sci U S A 1999; 96(15): 8745–8750.

4. Ryves WJ, Harwood AJ. Lithium inhibits glycogen synthase kinase-3 by competition for magnesium. Biochem Biophys Res Commun 2001; 280(3): 720–725.

5. Liang MH, Wendland JR, Chuang DM. Lithium inhibits Smad3/4 transactivation via increased CREB activity induced by enhanced PKA and AKT signaling. Mol Cell Neurosci 2008; 37(3): 440–453.

6. O’Brien WT, Harper AD, Jove F, Woodgett JR, Maretto S, Piccolo S et al. Glycogen synthase kinase-3beta haploinsufficiency mimics the behavioral and molecular effects of lithium. J Neurosci 2004; 24(30): 6791–6798.

7. O’Brien WT, Huang J, Buccafusca R, Garskof J, Valvezan AJ, Berry GT et al. Glycogen synthase kinase-3 is essential for beta-arrestin-2 complex formation and lithium-sensitive behaviors in mice. J Clin Invest 2011; 121(9): 3756–3762.

8. Hallcher LM, Sherman WR. The effects of lithium ion and other agents on the activity of myo-inositol-1-phosphatase from bovine brain. J Biol Chem 1980; 255(22): 10896–10901.

9. Sherman WR, Leavitt AL, Honchar MP, Hallcher LM, Phillips BE. Evidence that lithium alters phosphoinositide metabolism: chronic administration elevates primarily D-myo-inositol-1-phosphate in cerebral cortex of the rat. J Neurochem 1981; 36(6): 1947–1951.

10. Berridge MJ. Inositol trisphosphate and diacylglycerol: two interacting second messengers. Annu Rev Biochem 1987; 56: 159–193.

11. Berridge MJ, Downes CP, Hanley MR. Neural and developmental actions of lithium: a unifying hypothesis. Cell 1989; 59(3): 411–419.

12. Yuan P, Chen G, Manji HK. Lithium activates the c-Jun NH2-terminal kinases in vitro and in the CNS in vivo. J Neurochem 1999; 73(6): 2299–2309.

13. Casebolt TL, Jope RS. Effects of chronic lithium treatment on protein kinase C and cyclic AMP-dependent protein phosphorylation. Biol Psychiatry 1991; 29(3): 233–243.

14. Khayachi A, Abuzgaya M, Liu Y, Jiao C, Dejgaard K, Schorova L et al. Akt and AMPK activators rescue hyperexcitability in neurons from patients with bipolar disorder. EBioMedicine 2024; 104: 105161.

15. Stephenson EH, Higgins JMG. Pharmacological approaches to understanding protein kinase signaling networks. Front Pharmacol 2023; 14: 1310135.

16. Bruning F, Noya SB, Bange T, Koutsouli S, Rudolph JD, Tyagarajan SK et al. Sleep-wake cycles drive daily dynamics of synaptic phosphorylation. Science 2019; 366(6462).

17. Hastie CJ, McLauchlan HJ, Cohen P. Assay of protein kinases using radiolabeled ATP: a protocol. Nat Protoc 2006; 1(2): 968–971.

18. Gould TD, Chen G, Manji HK. In vivo evidence in the brain for lithium inhibition of glycogen synthase kinase-3. Neuropsychopharmacology 2004; 29(1): 32–38.

19. van Enkhuizen J, Milienne-Petiot M, Geyer MA, Young JW. Modeling bipolar disorder in mice by increasing acetylcholine or dopamine: chronic lithium treats most, but not all features. Psychopharmacology (Berl) 2015; 232(18): 3455–3467.

20. Can A, Blackwell RA, Piantadosi SC, Dao DT, O’Donnell KC, Gould TD. Antidepressant-like responses to lithium in genetically diverse mouse strains. Genes Brain Behav 2011; 10(4): 434–443.

21. Jensen J, Thomsen K, Olesen OV. Current determination of lithium-induced minimum sodium requirement in rats. Psychopharmacologia 1976; 45(3): 295–299.

22. Dunkley PR, Jarvie PE, Robinson PJ. A rapid Percoll gradient procedure for preparation of synaptosomes. Nat Protoc 2008; 3(11): 1718–1728.

23. Yang H, Smith P, Ma Y, Southworth E, Gopala Krishna V, Salerno B et al. Pervasive phenotypic effects of FBXO42 are promoted by regulation of PP4 phosphatase. EMBO J 2026; 45(4): 1332–1361.

24. Leeming MG, O’Callaghan S, Licata L, Iannuccelli M, Lo Surdo P, Micarelli E et al. Phosphomatics: interactive interrogation of substrate-kinase networks in global phosphoproteomics datasets. Bioinformatics 2021; 37(11): 1635–1636.

25. Buniello A, Suveges D, Cruz-Castillo C, Llinares MB, Cornu H, Lopez I et al. Open Targets Platform: facilitating therapeutic hypotheses building in drug discovery. Nucleic Acids Res 2025; 53(D1): D1467–D1475.

26. Kuleshov MV, Jones MR, Rouillard AD, Fernandez NF, Duan Q, Wang Z et al. Enrichr: a comprehensive gene set enrichment analysis web server 2016 update. Nucleic Acids Res 2016; 44(W1): W90–97.

27. Szklarczyk D, Kirsch R, Koutrouli M, Nastou K, Mehryary F, Hachilif R et al. The STRING database in 2023: protein-protein association networks and functional enrichment analyses for any sequenced genome of interest. Nucleic Acids Res 2023; 51(D1): D638–D646.

28. Otasek D, Morris JH, Boucas J, Pico AR, Demchak B. Cytoscape Automation: empowering workflow-based network analysis. Genome Biol 2019; 20(1): 185.

29. Eid S, Turk S, Volkamer A, Rippmann F, Fulle S. KinMap: a web-based tool for interactive navigation through human kinome data. BMC Bioinformatics 2017; 18(1): 16.

30. Karasawa T, Lombroso PJ. Disruption of striatal-enriched protein tyrosine phosphatase (STEP) function in neuropsychiatric disorders. Neurosci Res 2014; 89: 1–9.

31. Reichenstein M, Borovok N, Sheinin A, Brider T, Michaelevski I. Abelson Kinases Mediate the Depression of Spontaneous Synaptic Activity Induced by Amyloid Beta 1-42 Peptides. Cell Mol Neurobiol 2021; 41(3): 431–448.

32. Tanis KQ, Veach D, Duewel HS, Bornmann WG, Koleske AJ. Two distinct phosphorylation pathways have additive effects on Abl family kinase activation. Mol Cell Biol 2003; 23(11): 3884–3896.

33. Nagai T, Nakamuta S, Kuroda K, Nakauchi S, Nishioka T, Takano T et al. Phosphoproteomics of the Dopamine Pathway Enables Discovery of Rap1 Activation as a Reward Signal In Vivo. Neuron 2016; 89(3): 550–565.

34. Gurevich VV, Gurevich EV. GPCR Signaling Regulation: The Role of GRKs and Arrestins. Front Pharmacol 2019; 10: 125.

35. Johnson JL, Yaron TM, Huntsman EM, Kerelsky A, Song J, Regev A et al. An atlas of substrate specificities for the human serine/threonine kinome. Nature 2023; 613(7945): 759–766.

36. Yaron-Barir TM, Joughin BA, Huntsman EM, Kerelsky A, Cizin DM, Cohen BM et al. The intrinsic substrate specificity of the human tyrosine kinome. Nature 2024; 629(8014): 1174–1181.

37. Pitcher JA, Fredericks ZL, Stone WC, Premont RT, Stoffel RH, Koch WJ et al. Phosphatidylinositol 4,5-bisphosphate (PIP2)-enhanced G protein-coupled receptor kinase (GRK) activity. Location, structure, and regulation of the PIP2 binding site distinguishes the GRK subfamilies. J Biol Chem 1996; 271(40): 24907–24913.

38. Gainetdinov RR, Bohn LM, Sotnikova TD, Cyr M, Laakso A, Macrae AD et al. Dopaminergic supersensitivity in G protein-coupled receptor kinase 6-deficient mice. Neuron 2003; 38(2): 291–303.

39. Ashok AH, Marques TR, Jauhar S, Nour MM, Goodwin GM, Young AH et al. The dopamine hypothesis of bipolar affective disorder: the state of the art and implications for treatment. Mol Psychiatry 2017; 22(5): 666–679.

